# Modeling the effect of COVID-19 disease on the cardiac function: a computational study

**DOI:** 10.1101/2020.06.23.166421

**Authors:** Francesco Regazzoni, Christian Vergara, Luca Dede’, Paolo Zunino, Marco Guglielmo, Roberto Scrofani, Laura Fusini, Chiara Cogliati, Gianluca Pontone, Alfio Quarteroni

**Affiliations:** MOX, Dipartimento di Matematica, Politecnico di Milano, Milan, Italy; LABS, Dipartimento di Chimica, Materiali e Ingegneria Chimica, Politecnico di Milano, Milan, Italy; Centro Cardiologico Monzino IRCSS, Milan, Italy; Ospedale L. Sacco, Milan, Italy; Institute of Mathematics, Ecole Polytechnique Fédérale de Lausanne, Switzerland (*Professor Emeritus*)

**Author notes:** **ADDRESS FOR CORRESPONDENCE:** Francesco Regazzoni, Address: p.zza Leonardo da Vinci, 32, 20133 Milano.

**Keywords:** Covid-19, mathematical study, computational models, cardiac function, PV loop perturbation

## Abstract

**Background:** The effect of COVID-19 on the cardiac function and on the vascular system increases the morbidity and mortality of infected subjects with cardiovascular diseases.

**Objectives:** To provide preliminary results on cardiac global outcomes (such as cardiac output, ventricular pressures) obtained by means of computational models in plausible scenarios characterized by COVID-19.

**Methods:** We considered a lumped parameters computational model of the cardiovascular system, which models, from the mechanical point of view, the systemic and pulmonary circulations, the four cardiac valves and the four heart chambers, through mathematical equations of the underlying physical processes. To study the effect of COVID-19, we varied the heart rate, the contractility and the pulmonary resistances in suitable ranges.

**Results:** Our computations on individuals with both otherwise normal and impaired cardiac functions revealed that COVID-19 worsen cardiac function, as shown by a decrease of some cardiac biomarkers values such as cardiac output and ejection fraction. In the case of existing impaired cardiac function, the presence of COVID-19 lead to values outside the normal ranges.

**Conclusions:** Computational models revealed to be an effective tool to study the effect of COVID-19 on the cardiovascular system. Such effect could be significant for patients with impaired cardiac function. This is especially useful to perform a sensitivity analysis of the hemodynamics for different conditions.

**CONDENSED ABSTRACT:** Emerging studies address how COVID-19 infection might impact the cardiovascular system. This relates particularly to the development of myocardial injury, acute coronary syndrome, myocarditis, arrhythmia, and heart failure. Prospective treatment approach is advised for these patients. By the assessment of conventional important biomarkers obtained with new sources as a 0-dimentional computational model, we propose a new study protocol as an effective method to evaluate short-term prognosis. The clinical protocol proposed will help to rapidly identify which patients require intensive monitoring, diagnostic strategy and most adequate therapy.

## INTRODUCTION

The Coronavirus disease 2019 (COVID-19) caused by the severe acute respiratory syndrome coronavirus 2 (SARS-CoV-2) primarily affects the respiratory system, although others organ systems are also involved and in particular cardiovascular complications may occur.

Among patients with COVID-19, there is a high prevalence of previous history of cardiovascular diseases (CVD) and/or CVD risk factors and increased morbidity and mortality in those patients have been recently reported in several studies [M-JAMA-20, G-JAMA-20, R-ICM-20, Z-Lancet-20, L-JACC-20].

The virus invades cardiomyocytes by binding to angiotensin converting enzyme 2 on the cell surface, resulting in myocardial injury which elevates troponin I levels. In particular, the COVID-19 infection can lead to myocarditis, vascular inflammation, arrhythmias, acute heart failure, and in the most severe cases, cardiogenic shock and death [M-JAMA-20, G-JAMA-20]. Studies of patients with COVID-19 in China indicated significantly higher in-hospital mortality rate in patients who also have myocardial injury. Acute infections increase the risk of acute coronary syndrome. COVID-19 may increase circulating cytokines, which have the potential to cause instability and rupture of atherosclerotic plaques. Fever, stress, electrolyte disturbances and use of antiviral drugs in patients with COVID-19 may all have pro-arrhythmic effects. This may be an issue for patients with inherited arrhythmia syndromes, such as long QT syndrome, Brugada syndrome, short QT syndrome and catecholaminergic polymorphic ventricular tachycardia. For these patients, additional precautions and specialised management are advised.

Moreover, some studies highlighted that, for patients hospitalized with COVID-19, a higher rate of comorbidities including hypertension and coronary artery disease characterized fatal events when compared to survivors [Z-Lancet-20] and myocardial injury was prevalent [L-JACC-20]. In any case, the interplay of COVID-19 with cardiovascular complications both in healthy individuals and in patients with pre-existing CVD is still far from being fully understood. The mechanisms of cardiac injury in patients with COVID-19 are not well established but it is reasonable to identify both direct and indirect mechanisms responsible for cardiac failure.

Possible causes of the effect of COVID-19 on cardiac impairment are due to an increased pulmonary vascular resistance consistent with severe hyperinflammation and micro-thrombosis [M-CCM-20]. On the other side, the decreased blood saturation secondary to respiratory failure may hamper the cardiomyocytes contractility, possibly leading to a decreased cardiac output. Both as compensatory mechanism for the reduced oxygen concentration in blood and consequence of fever rise due to the COVID-19 inflammation, the heart rate may significantly increase.

In this study, we provided examples of possible scenarios that investigate the effect of COVID-19 on some cardiac features of interest and other vital parameters. With this aim, we exploited the predictive nature of *computational methods* [Q-CUP-20] that are based on the numerical solution (i.e. aided by a computer) of the mathematical equations underlying the physical processes under investigation. In this context, a *zero-dimensional* (0D, also known as lumped parameters) model that simulates systemic and pulmonary blood dynamics together with the cardiac function has been considered [O-ABE-00]. This tool can account for a broad range of the model parameters, to simulate variations in pulmonary resistances, heart rate, and cardiomyocyte contractility in accordance with COVID-19 infection, and to compute the corresponding changes of some meaningful outputs such as stroke volume, ejection fraction, cardiac output, arterial and pulmonary pressures.

The use of a 0D mathematical model allowed us to better understand the strong interplay between local changes (such as the reduced contractility), global variables (such as peripheral resistances, heart rate) and cardiac function. This is of utmost importance in view of determining the possible effects of COVID-19 infection on patients affected by CVD and, on the other hand, to better evaluate the impact of CVD on COVID-19. A possible scenario of COVID-19 infection in an individual exhibiting a reduced ejection fraction and cardiac output is investigated to sustain the power of our computational study and its possible use for patient-specific studies.

## METHODS

### The zero-dimensional computational model

A 0D model provides a mathematical description of the function of several compartments in the cardiovascular system; their number and locations depend on the complexity of the model at hand and its level of detail. The model consists in a system of differential equations that translate physical principles such as conservation of mass and momentum. Its solution provides the values of flow rates and pressures for each compartment [R-CVS-01]. This system can be obtained by exploiting the electric analogy, where the current represents the blood flow, the voltage is the pressure, the resistance represents the blood resistance, the conductance the vessel compliance, and the inductance the blood inertia.

Referring to Figure 1, our 0D model was composed by:

1. The four heart chambers, whose mechanical behavior was characterized by suitable model parameters. For example, for the left ventricle (LV) the unloaded volumes (i.e. volume at zero-pressure) V0_LV_ and the time-varying elastances E_LV_(t)=EB_LV_ + EA_LV_ * f_LV_ (t) were considered, where EB_LV_ is the passive elastance (i.e the inverse of the compliance), EA_LV_ the maximum active elastance, and f_LV_ a function of time ranging values between 0 and 1 that accounts for the activation phases. Similar parameters definitions were introduced for the right ventricle (RV), left atrium (LA), and right atrium (RA);
2. The four cardiac valves modeled as diodes;
3. The systemic circulation, characterized by the arterial resistance, compliance and inductance R_AR_^SYS^, C_AR_^SYS^, L_AR_^SYS^, respectively, and by the same quantities related to the venous system, i.e. R_VEN_^SYS^, C_VEN_ ^SYS^, L_VEN_ ^SYS^;
4. The pulmonary circulation, characterized as above by the systemic parameters R_AR_^PUL^, C_AR_^PUL^, L_AR_^PUL^, and by the venous ones R_VEN_^PUL^, C_VEN_^PUL^, L_VEN_^PUL^.

**Central illustration - Figure 1.**
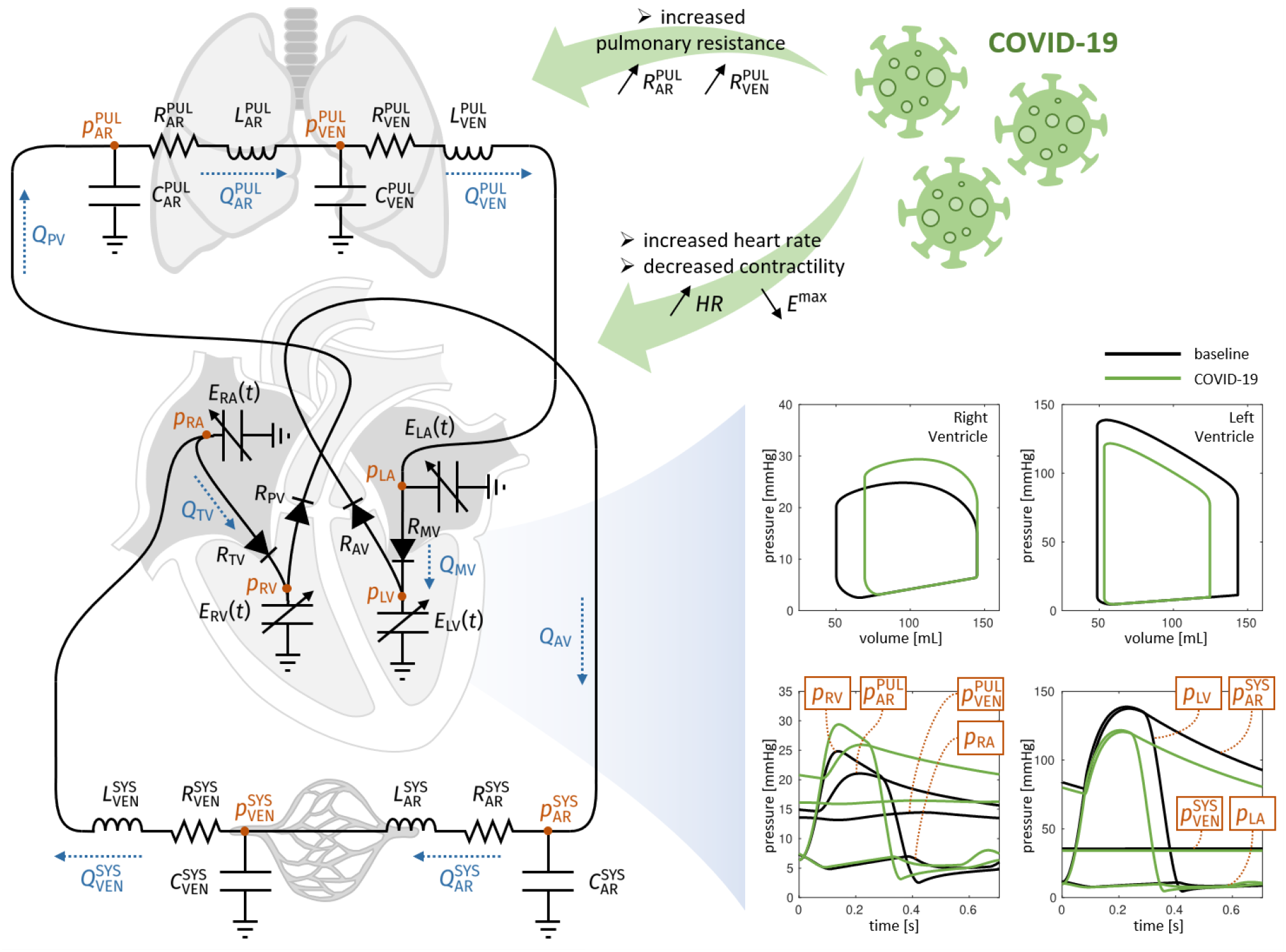
0D model, schematic inputs, and representative outcomes. 0D model (on the left), effect of COVID-19 on the input parameters (up, right), and representative outcomes in terms of PV loops and pressure behaviors (bottom, right). p_LV_ = left ventricle pressure, p_LA_ = left atrium pressure, p_RV_ = right ventricle pressure, p_RA_ = right atrium pressure, p_AR_^SYS^ = systemic arterial pressure, p_VEN_^SYS^ = systemic venous pressure, p_AR_^PUL^ = pulmonary arterial pressure, p_VEN_^PUL^ = pulmonary venous pressure

For a recent study on a 0D model accounting for the pulmonary circulation, see also [F-IJNMBE-20].

### Variation of the model parameters

The quantities mentioned in the previous list are the parameters of the model and, together with the heart rate (HR), they fully characterize the individual or patient’s condition (healthy or non-healthy). The 0D model thus provides a tool that is able to compute flow rate and pressure at each compartment given a chosen set of parameters. With this aim, we identified three sources of changes in the above-mentioned parameters and corresponding reasonable value ranges:

1. HR: Clinical evidences showed that tachycardia was very common in COVID-19 patients [X-EHJ-20]. Accordingly, we placed the values of HR in the range [60,100] bpm;
2. Wood number (W): It is defined as

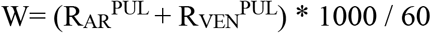

and represents an information about the pulmonary resistances. Since, according to clinical evidences [Q-CC-20], pulmonary resistances (and thus W) increase in presence of COVID-19, we placed the value of W in the range between 1.13 (healthy) and 3.39. More precisely, we multiplied the baseline value R_VEN_^PUL^_-BASE_ (see Table 1) by a factor k_R_ in the range [1,3]:

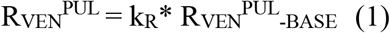

**Table 1.** Baseline values of the parameters used in the 0D model.

| Parameter | Values | Unit |
| --- | --- | --- |
| HR | 72 | bpm |
| $EB_{LV}$ | 0.08 | mmHg / mL |
| $EB_{RV}$ | 0.05 | mmHg / mL |
| $EB_{LA}$ | 0.09 | mmHg / mL |
| $EB_{RA}$ | 0.07 | mmHg / mL |
| $EA_{LV-bsln}$ | 2.75 | mmHg / mL |
| $EA_{RV-bsln}$ | 0.55 | mmHg / mL |
| $EA_{LA-bsln}$ | 0.07 | mmHg / mL |
| $EA_{RA-bsln}$ | 0.06 | mmHg / mL |
| $R_{AR}^{SYS}$ | 0.64 | mmHg s / mL |
| $C_{AR}^{SYS}$ | 1.2 | mL / mmHg |
| $L_{AR}^{SYS}$ | 0.005 | mmHg s<sup>2</sup> / mL |
| $R_{VEN}^{SYS}$ | 0.26 | mmHg s / mL |
| $C_{VEN}^{SYS}$ | 60 | mL / mmHg |
| $L_{VEN}^{SYS}$ | 0.0005 | mmHg s<sup>2</sup> / mL |
| $R_{AR}^{PUL-bsln}$ | 0.0321 | mmHg s / mL |
| $C_{AR}^{PUL}$ | 10 | mL / mmHg |
| $L_{AR}^{PUL}$ | 0.0005 | mmHg s<sup>2</sup> / mL |
| $R_{VEN}^{PUL-bsln}$ | 0.0357 | mmHg s / mL |
| $C_{VEN}^{PUL}$ | 16 | mL / mmHg |
| $L_{VEN}^{PUL}$ | 0.0005 | mmHg s<sup>2</sup> / mL |

We also assumed that R_AR_^PUL^/R_VEN_^PUL^ = 9/10 [H-IJNMBE-17]. Analogous definitions hold true for the systemic resistances;

EA_XX_: Clinical evidences showed that COVID-19 may produce a decreased blood saturation and consequently also a decreased cardiomyocyte contractility. To account for this, we decreased the value of the four active elastance EA_XX_ by multiplying the baseline values EA_XX-BASE_ (see Table 1) by a percentage factor k_EA_ in the range [50,100]% [W-CR-88,B-PLOS-18]:

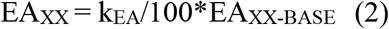

## RESULTS

First, we considered the setting reported in Table 1, corresponding to a patient in healthy conditions, henceforth denoted as *baseline* (bsln). In Figure 2 we reported the outcomes predicted by the 0D model for this baseline setting. Our computational model allowed to reproduce the ventricle pressure-volume (PV) loops as well as the evolution in time of volumes and pressures. Table 2 reports a list of cardiac measurements computed by the baseline simulation. In particular, we found Cardiac Output (CO) = 6.8 L/min, Stroke Volume (SV) = 95 mL, Ejection Fraction (EF) = 66% for both LV and RV, thus the simulation reflected an overall normal cardiac function. Also, LV and RV were characterized by systolic and end-diastolic volumes within the control ranges (LV End Diastolic Volume (EDV) = 144 mL, LV End Systolic Volume (ESV) = 49 mL, RV EDV = 145 mL, RV ESV = 50 mL) [M-EHJ-06, M-JCMR-06, C-BMC-09].

**Figure 2.**
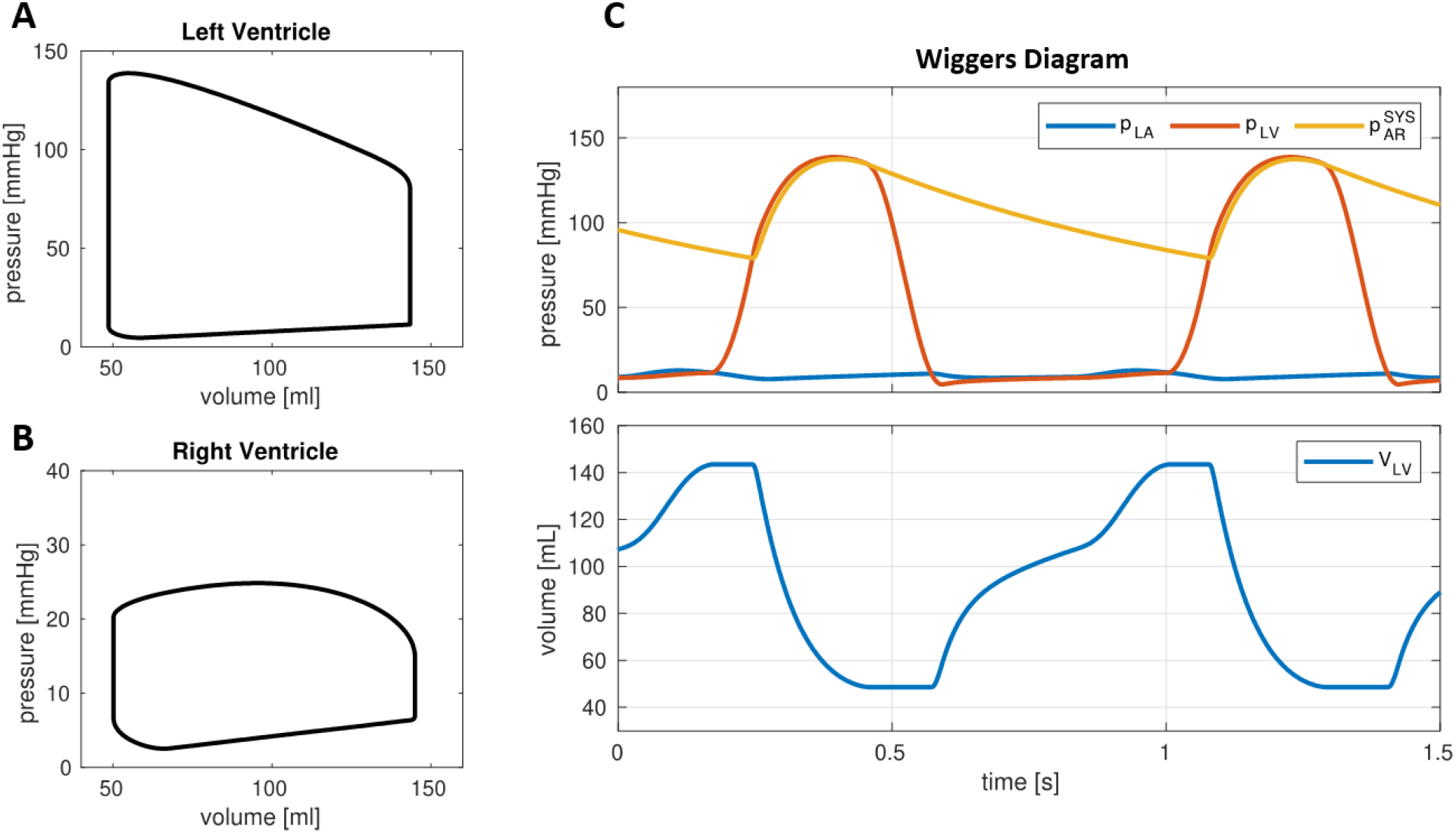
PV loops and pressure for the baseline case. Output of the 0D model resulting from the baseline parameters. A: Left ventricle PV loop. B: Right ventricle PV loop. C: Wiggers diagram, showing the evolution, for two heartbeats, of the left ventricle pressure (p_LV_) and volume (V_LV_), the left atrium pressure (p_LA_) and the systemic arterial circulation pressure (p_AR_^SYS^), which is representative of the aortic pressure

**Table 2.** Model parameters (inputs) for simulating the effects of mild (Test (1a)) and severe (Test (1b)) COVID-19 infections in an individual with otherwise normal cardiac function (baseline), and computed outputs. For the pulmonary resistances and maximum active elastances current values are derived by the baseline ones owing to equations (1) and (2). Pictures of computed outputs in Figure 3A.

| Parameter | Unit | bsln | Test (1a) | Test (1b) |
| --- | --- | --- | --- | --- |
| **MODEL INPUTS** |  |  |  |  |
| Contractility wrt bsln ($k_{EA}$) | % | 100 | 85 | 70 |
| HR | bpm | 72 | 86 | 100 |
| Systemic resistance | Wood | 15 | 15 | 15 |
| Pulmonary resistance | Wood | 1.13 ($k_R=1$) | 2.26 ($k_R=2$) | 3.39 ($k_R=3$) |
| **MODEL OUTPUTS** |  |  |  |  |
| **General** |  |  |  |  |
| CO | L/min | 6.8 | 6.4 | 5.9 |
| SV | mL | 95 | 75 | 59 |
| **Left Ventricle** |  |  |  |  |
| LV EDV | mL | 144 | 127 | 117 |
| LV ESV | mL | 49 | 52 | 58 |
| LV EF | % | 66 | 59 | 51 |
| LV Pmax | mmHg | 139 | 127 | 116 |
| LV dP/dt max | mmHg/s | 1649 | 1658 | 1579 |
| **Right Ventricle** |  |  |  |  |
| RV EDV | mL | 145 | 142 | 148 |
| RV ESV | mL | 50 | 67 | 89 |
| RV EF | % | 66 | 53 | 40 |
| RV Pmax | mmHg | 25 | 30 | 35 |
| RV dP/dt max | mmHg/s | 256 | 338 | 383 |
| **Systemic circulation** |  |  |  |  |
| PA min | mmHg | 79 | 79 | 76 |
| PA max | mmHg | 137 | 126 | 115 |
| **Pulmonary circulation** |  |  |  |  |
| PAP mean | mmHg | 17 | 23 | 29 |

With the baseline setting as a starting point, we investigated the effect of the variations mentioned in the Methods section, to study a possible impact of COVID-19 on the cardiac function. In particular, we considered two settings, corresponding to a mild and to a more severe COVID-19 infection, that we denoted as Test (1a) and Test (1b), respectively. In Test (1a), to reflect the effects of a reduced blood saturation due to the impaired pulmonary function, we decreased the contractility of the whole heart muscle to the 85% of the baseline value. Moreover, we increased HR from 72 bpm to 86 bpm and we increased the pulmonary resistance by a factor 2 (this corresponds to a Wood number 2.26). In Test (1b), instead, we accentuated the three abovementioned effects, with a contractility of 70% of the baseline, a HR of 100 bpm and an increase of the pulmonary resistances of a factor 3 (Wood number 3.39). The PV loops corresponding to the settings (1a) and (1b) are shown in Figure 3A, while Table 2 reports the corresponding cardiac measurements. These tests showed that the LV PV loops was impacted by the hypoxemia that reduced their contractility. Overall, compared to baseline values, the reduced cardiac contractility provides a decrease in SV of 21% for Test (1a) and 38% for Test (1b), together with a reduction in LV EF of 11% and 25% for Test (1a) and Test (1b), respectively. This effect was partially compensated by the increase of HR, which can be considered – in hypoxic patients – as a self-regulatory mechanism aimed to preserve the CO [B-PLOS-18]. As a matter of fact, the CO decreased of 6% from baseline for Test (1a) and of 13% for Test (1b). Besides the LV function, also the RV function is significantly affected by COVID-19 related variations of the circulation model. Specifically, RV EF decreased of 20% (Test (1a)) and 39% (Test (1b)), thus in percentage more than for LV. Moreover, the increased pulmonary resistance induced a remarkable growth of the RV maximal pressure (20% and 40% for Test (1a) and (1b), respectively) and the RV ESV reports a significant increase (34% and 78%).

**Figure 3.**
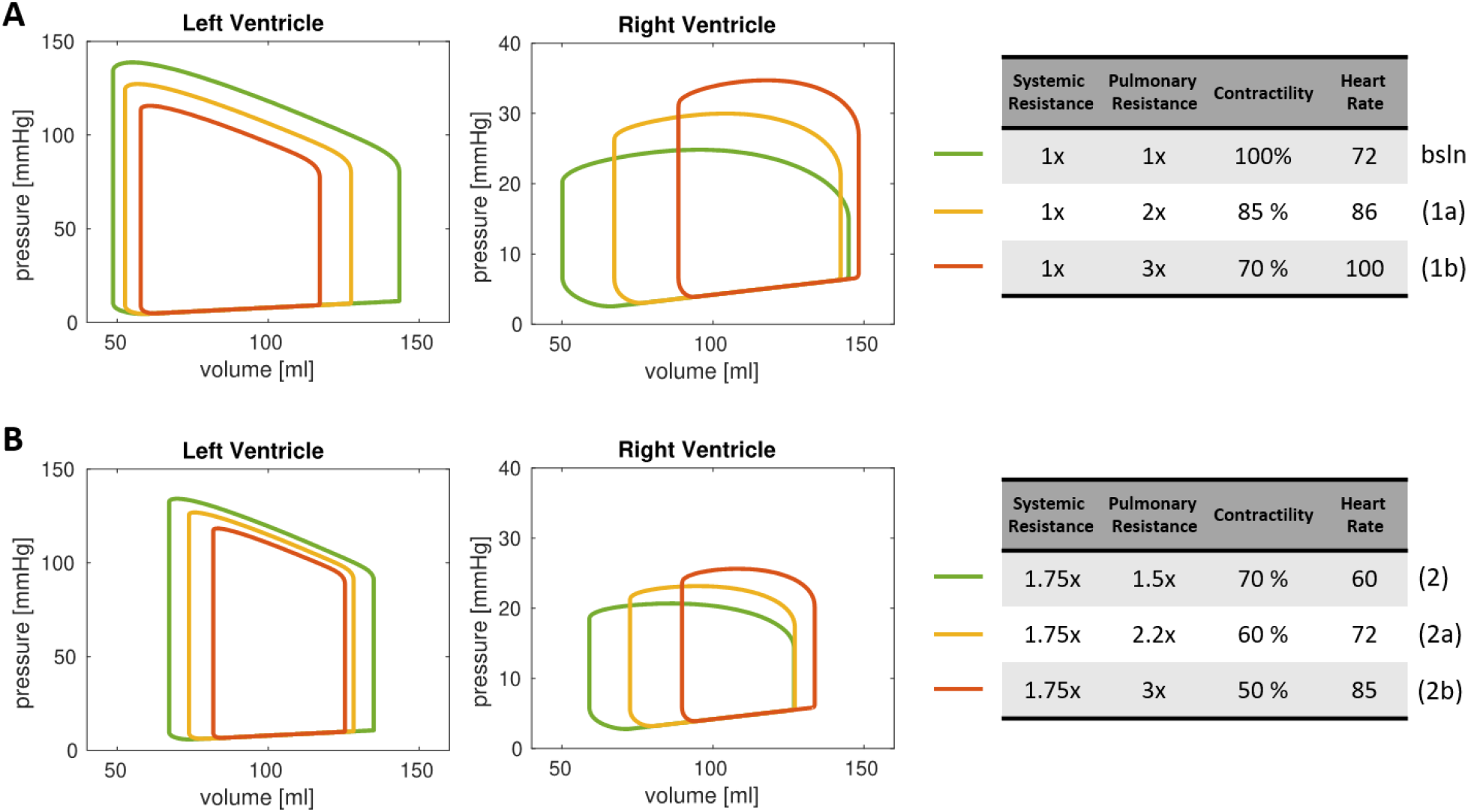
PV loops for the COVID-19 scenarios. PV loops of LV and RV for different scenarios of COVID-19 infections. A: results obtained with model parameters simulating the effects of mild (1a) and severe (1b) COVID-19 infections in an individual with otherwise normal cardiac function (baseline). B: results obtained by model parameters simulating the effects of mild (2a) and severe (2b) COVID-19 infections in an individual with impaired cardiac function.

We further exploited the predictive power of our computational model by simulating the effects of COVID-19 infection for an individual with mildly impaired LV cardiac function. In particular, we assumed that for this subject, in absence of COVID-19, contractility was decreased to 70% of baseline, HR equal to 60 bpm, pulmonary resistance equal to 1.70 Wood (x1.5 of baseline) and systemic resistance equal to 26.25 (x1.75 of baseline) (Test (2), see Table 3). The outputs computed by the 0D model were: CO = 4.1 L/min, LV SV = 68 mL, LV EF = 50% and RV EF = 53% ; furtherly, we found LV EDV = 135 mL, LV ESV = 67 mL, RV EDV = 127 mL, RV ESV = 59 mL, that however were still in the normal ranges [M-EHJ-06, M-JCMR-06, C-BMC-09].

**Table 3.** Model parameters for simulating the effects of mild (Test (2b)) and severe (Test (2c)) COVID-19 infections in an individual with impaired cardiac function (Test (2a)), and computed outputs. For the pulmonary resistances and maximum active elastances current values are derived by the baseline ones owing to equations (1) and (2). Baseline: k_R_ =1, k_EA_ =100. Pictures of computed output in Figure 3B

| Parameter | Unit | Test (2a) | Test (2b) | Test (2c) |
| --- | --- | --- | --- | --- |
| **MODEL INPUTS** |  |  |  |  |
| Contractility wrt bsln ($k_{EA}$) | % | 70 | 60 | 50 |
| HR | bpm | 60 | 72 | 85 |
| Systemic resistance | Wood | 26.25 | 26.25 | 26.25 |
| Pulmonary resistance | Wood | 1.70 ($k_R=1.5$) | 2.54 ($k_R=2.25$) | 3.39 ($k_R=3$) |
| **MODEL OUTPUTS** |  |  |  |  |
| **General** |  |  |  |  |
| CO | L/min | 4.1 | 3.9 | 3.7 |
| SV | mL | 68 | 55 | 44 |
| EF | % | 50 | 43 | 35 |
| **Left Ventricle** |  |  |  |  |
| LV EDV | mL | 135 | 128 | 125 |
| LV ESV | mL | 67 | 73 | 82 |
| LV EF | % | 50 | 43 | 35 |
| LV Pmax | mmHg | 134 | 127 | 117 |
| LV dP/dt max | mmHg/s | 1100 | 1131 | 1115 |
| **Right Ventricle** |  |  |  |  |
| RV EDV | mL | 127 | 127 | 134 |
| RV ESV | mL | 59 | 72 | 90 |
| RV EF | % | 53 | 43 | 33 |
| RV Pmax | mmHg | 21 | 23 | 26 |
| RV dP/dt max | mmHg/s | 159 | 189 | 210 |
| **Systemic circulation** |  |  |  |  |
| DBP | mmHg | 90 | 90 | 88 |
| SBP | mmHg | 133 | 126 | 118 |
| **Pulmonary circulation** |  |  |  |  |
| PAP mean | mmHg | 16 | 19 | 22 |

As in Test (1), we considered two scenarios of COVID-19 infections by furtherly decreasing the heart contractility (to 60% and 50% of baseline), increasing the HR (to 72 and 85 bpm) and increasing the pulmonary resistance (to 2.54 and 3.39 Wood) for Tests (2a) and (2b), respectively. The data and results are reported in Figure 3B and Table 3. The outputs of our numerical simulations highlighted that, in an individual with impaired LV cardiac function, the COVID-19 infection furtherly worsen the cardiac functional parameters of both chambers. In particular, we found a reduction in LV EF of 14% and 30% for Test (2a) and Test (2b), respectively, in LV EF of 14% and 30%, in RV EF of 19% and 38%, and in CO of 5% and 10%.

## DISCUSSION

### Discussion of the results

Our computational model is able to predict possible effects of COVID-19 infection on the cardiocirculatory system, and particularly on the cardiac function. Our study confirmed that the infection is able to impact both the left and right hearts, by decreasing EF and CO, whose decrements are only partially compensated by the HR increase that typically is shown in COVID-19 patients. Maximum LV pressure was reduced, while maximum RV pressure was increased, albeit the latter remains within the normal range of values. Our computational model predicted that the right heart is strongly linked, from the physiopathological standpoint, to the pulmonary function. Indeed, the increased pulmonary resistance significantly affects both the RV pressure and the RV EF.

These results seem to be well in keeping with recent data reporting that lower RV systolic function echocardiographic parameters, although still in the normal range, are more frequent in Covid patients with poor prognosis, thus suggesting that the absence of an increase in contractility in response to an increase in pulmonary resistance/pressures could play a role in the evolution of the illness [L-JACC-20].

As the respiratory failure worsen, RV function is supposed to be more susceptible to impairment due to increased RV afterload and in fact we have reported a progressive decrease in RV EF volume from Test (1a) to Test (1b) for a previously healthy subject. Since recent reports [M-JAMA-20, G-JAMA-20, R-ICM-20, Z-Lancet-20, L-JACC-20] suggest that CVD is a common finding in patients with COVID-19 and is associated with a worse prognosis, we also simulated the cardiac function and its response to COVID-19 in a patient with history of cardiac dysfunction. Although we measured a reduction in LV EF when we simulate a worse COVID-19 infection, we measured a more severe reduction in RV EF compared to the reduction in LV EF and this finding could explain the worse outcome in case of previous CVD. In fact, RV dysfunction is known to be an important determinant of symptoms and a powerful marker of poor prognosis in patients with chronic heart failure [G-EJHF-17, C-CCI-18].

To predict the effect of the reduced contractility, we estimated the End Systolic Pressure Volume Relationship (ESPVR) as the slope of the PV loop curve in the upper left corner. As a matter of fact, as highlighted by Figures 3A and 3B, left, the depressed inotropy reflected in a decreased ESPVR. This effect has been reported in several experiments as a direct consequence of the reduced oxygen blood content [W-CR-88, B-PLOS-18]. Moreover, the LV function experienced a reduced systolic pressure and a reduced preload, thus resulting in a significant decrease of the PVA (pressure-volume area), a further biomarker that is known to correlate with hypoxia [K-AJP-65, B-PLOS-18].

From our preliminary study, we highlighted that COVID-19 infection in a patient with healthy cardiac function may be able to produce alterations that maintain biomarkers within the normal range of values. Conversely, starting from a scenario of an individual with impaired cardiac function, where however the values of biomarkers were within normal ranges [M-EHJ-06, M-JCMR-06, C-BMC-09], our simulations revealed that the COVID-19 infection could further worsen the cardiac biomarkers, specifically by further reducing CO and SV below the normal values, other than severely impacting the LV and RV EFs.

The computational model has been proposed to carry out sensitivity analyses in a very rapid, reliable and reproducible way. It compares the effect of very complex clinical situation of the patient employed in the medical practice on the outcome of treatments in several relevant cases.

### Study limitations

This study was based on a 0D computational model, which considers compartmental flow rates and pressures. A more detailed study for predicting pointwise quantities of interest in the heart could be carried out by means of a 3D-0D model, where the heart (or a part of it) is simulated by means of a 3D electromechanical model, whose geometry can be personalized starting from clinical images (CT or MRI) acquired from a specific patient. This is currently under study (Regazzoni F, Salvador M, Africa P, Fedele M, Dede’ L, Quarteroni A. Electromechanical modeling of the human heart coupled with a circulation model of the whole cardiovascular system, in preparation).

Our computational model neglected the effect of pressure variations due to respiration on the rib cage, hence on the cardiocirculatory system. Albeit we deem this assumption to have a very limited impact on the outcomes of this study, we plan to improve our 0D model for further studies on this topic.

## CONCLUSIONS

Computational models could be an effective tool to study the effect of COVID-19 on the cardiac function. Our preliminary 0D model computations revealed a possible worsening of SV, CO, LV EF and RV EF and other biomarkers both for individuals with otherwise normal cardiac function and with impaired cardiac function. In the latter case, these values were below normal values even if they were within normal ranges in the case without COVID-19 effects. This provided a quantitative, although speculative, confirmation that COVID-19 could have important consequences on the cardiac function.

## ACKNOWLEDGMENTS

C. Vergara has been partially supported by the H2020-MSCA-ITN-2017, EU project 765374 “ROMSOC - Reduced Order Modelling, Simulation and Optimization of Coupled systems”. L. Dede’, A. Quarteroni, C. Vergara and P. Zunino have been partially supported by the Italian research project MIUR PRIN17 2017AXL54F “Modeling the heart across the scales: from cardiac cells to the whole organ”.

## ABBREVIATION LIST

HR: Heart Rate
EDV: End Diastolic Volume
ESV: End Systolic Volume
CO: Cardiac Output
SV: Stroke Volume
EF: Ejection Fraction
DBP: (Systemic) Diastolic Blood Pressure
SBP: (Systemic) Systolic Blood Pressure
PAP: Pulmonary artery pressure

## Notes

**FUNDING**: This project has received funding from the European Research Council (ERC) under the European Union Horizon 2020 research and innovation programme (grant agreement No 740132, iHEART - An Integrated Heart Model for the simulation of the cardiac function, P.I. Prof. A. Quarteroni).

**DISCLOSURES**: The authors declare that they have no conflict of interest with industries or other.

### Competing Interest Statement

The authors have declared no competing interest.

